# Supercharged Assembly: A Broad-Spectrum Mechanism of Action for Drugs that Undermine Controlled HIV-1 Viral Capsid Formation

**DOI:** 10.1101/565515

**Authors:** Alexander J. Pak, John M. A. Grime, Alvin Yu, Gregory A. Voth

## Abstract

The early and late stages of human immunodeficiency virus (HIV) replication are orchestrated by the capsid (CA) protein, which self-assembles into a conical protein shell during viral maturation. Small molecule drugs known as capsid inhibitors (CIs) impede the highly-regulated activity of CA. Intriguingly, a few CIs, such as PF-3450074 (PF74) and GS-CA1, exhibit effects at multiple stages of the viral lifecycle at effective concentrations in the pM to nM regimes, while the majority of CIs target a single stage of the viral lifecycle and are effective at nM to µM concentrations. In this work, we use coarse-grained (CG) molecular dynamics simulations to elucidate the molecular mechanisms that enable CIs to have such curious broad-spectrum activity. Our quantitatively analyzed findings show that CIs can have a profound impact on the hierarchical self-assembly of CA by perturbing the population of small CA oligomers. The self-assembly process is accelerated by the emergence of alternative assembly pathways that favor the rapid incorporation of CA pentamers, and leads to increased structural pleomorphism of mature capsids. Two relevant phenotypes are observed: (1) eccentric capsid formation that may fail to encase the viral genome and (2) rapid disassembly of the capsid, which express at late and early stages of infection, respectively. Finally, our study emphasizes the importance of adopting a dynamical perspective on inhibitory mechanisms and provides a basis for the design of future therapeutics that are effective at low stoichiometric ratios of drug to protein.

## Introduction

Over the last few decades, antiretroviral therapy (ART) for human immunodeficiency virus type 1 (HIV-1) has made substantial progress.^1–3^ These advances can be attributed, in part, to the identification of multiple enzymes that are critical to the HIV-1 lifecycle,^4–6^ such as reverse transcriptase, integrase, and protease, as well as the development of small molecules that competitively inhibit their activities. Current state-of-the-art ARTs consist of a combination (cART) approach that utilizes multiple drugs, each active against different targets; this strategy is also referred to as highly-active ART (HAART). Nonetheless, the primary challenges for HAARTs are to maintain efficacy, safety, and tolerability.^1–2^ In particular, individual genetic variation has been responsible for drug resistance in patients.^7^ Treatment regimens must therefore adapt to these circumstances, e.g., through the introduction of drugs with alternative mechanisms of action, since clinically available options are limited.

An emerging class of drugs aims to disrupt the activity of the Gag (group specific antigen) polyprotein, which is responsible for coordinating the late stages of the viral lifecycle.^8–9^ The capsid (CA) domain of Gag is an attractive therapeutic target, since both the assembly and maturation of infectious viral particles are mediated by interactions between CA protein domains.^10–16^ During maturation, for example, CA self-assembles into a conical capsid (i.e., the mature core), composed of more than one thousand CA monomers, that encases the viral genome.^17–18^ Given the functional significance of the CA domain, it is also important to note that its sequence is highly conserved (around 70%) amongst HIV-1 subtypes, thereby reducing the risk of viral polymorphism.^19–20^ Drugs that target CA are known as capsid inhibitors (CIs) and have been studied for nearly a decade.^21–22^ Several candidates have been identified that demonstrate the feasibility of the CI approach, such as Bevirimat and PF-3450074 (PF74), which have half maximal effective concentrations (EC_50_), a measure of drug potencies, at nM to µM concentrations.^23–24^ Most recently, GS-CA1 was introduced as a promising CI with an EC_50_ around 85 pM concentration, and targets the same binding pocket as PF74 during early and late stages of viral infection.^25^ Nonetheless, a clinically viable CI has yet to appear. A molecular understanding of the mechanism of action for CIs is currently lacking, and represents a barrier for the development of new therapeutics.

Perhaps the most extensively studied CI, PF74 serves as a useful example to highlight the complexity of potential mechanisms for CIs. Initial reports of PF74 have suggested that capsid destabilization is the primary mechanism of action for the drug.^23^ Further capsid disassembly assays have shown that PF74 induces viral uncoating,^26–28^ although a contradictory study, using assays at similar concentrations of PF74, observed no discernible effects on viral uncoating and reverse transcription.^29^ Interestingly, similar assays that were performed on preassembled CA tubules have shown that PF74 has a stabilization effect.^28, 30–31^ Several binding assays have indicated that PF74 preferentially binds to CA multimers rather than isolated CA,^23, 28, 31–32^ and have found that CA assembly rates increase with PF74 present.^23, 31^ One hypothesis that may resolve the aforementioned contradiction is that PF74 affects the kinetics of the CA assembly process, which warrants further investigation. Another hypothesis has also been presented on the basis of the crystal structure of PF74 in complex with CA hexamers. Since the drug binds to a pocket at the interface between the N-terminal domain (NTD) and C-terminal domain (CTD) of adjacent CA monomers,^23, 28^ PF74 binding may preclude variable curvatures in CA oligomers that must be adopted for closed capsids. However, since this binding pocket has also been associated with cleavage and polyadenylation specific factor 6 (CPSF6) and nucleoporin 153 kDa (NUP153), two cellular transport factors that aid nuclear import,^33–34^ another possibility is that nuclear integration of the virus is abrogated due to competitive inhibition by PF74.^29^ Taken together, these observations suggest that PF74 (and related CIs, such as GS-CA1) can exhibit complex, multimodal mechanisms of action that target viral maturation, uncoating, reverse transcription, and nuclear import. Understanding the molecular mechanisms by which CIs produce one key aspect (capsid assembly/disassembly) of such broad activity is the focus of this work.

We use coarse-grained (CG) molecular dynamics simulations based on prior CG models^11^ to investigate the impact of CIs on viral infectivity. We hypothesize that the role of CIs is to perturb the population of small, intermediate oligomers of CA. These effects are implicitly incorporated into our CG model and used to examine the subsequent implications on late and early stages of the viral lifecycle. We simulate both capsid assembly and disassembly and emphasize the differences between the assembly/disassembly pathways in models for both apo and CI-bound capsids. Our findings suggest that CIs have multiple modes of action by increasing the population of mature capsids that either (i) fail to enclose viral RNA during maturation or (ii) spontaneously disassemble before transport and nuclear integration. Furthermore, our study implies that the targeting of CA oligomers, which requires low stoichiometric loadings of drugs to CA, is a possible design strategy for future therapeutics.

## Results and Discussion

We begin by considering the hierarchical nature of viral capsid assembly, which is schematically shown in Fig. 1. The capsid is an enclosed protein core that is composed of tiled CA hexamers with 12 CA pentamers incorporated for topological completeness.^17–18^ In the cytoplasm, CA exists in a dynamic equilibrium between CA monomers and dimers.^35^ Furthermore, the conformation of CA dimers is inherently dynamic, as depicted in the top row of Fig. 1, and importantly, only adopts conformations compatible with the mature capsid with 5-10% probability (i.e., the conformer observed in mature CA hexamers, which is shown in the upper half of the red box in Fig. 1).^36^ CA dimers associate during the assembly process to form the final capsid structure. Prior CG simulations^10–11,37^and a kinetic model^38^ have shown that a key intermediate is the trimer of dimers (TOD) structure, depicted in the lower half of the red box in Fig. 1. Three TODs form a complete CA hexamer that can simultaneously template adjacent hexamers. As CA oligomers increase in size, there are numerous possible assembly outcomes, including stalled or malformed assemblies. However, under the precise conditions of weak CA-CA interactions with highly-specific association interfaces, which are only accessible in a limited manner, the self-assembly process expresses the characteristic fullerene core morphology.^11^ These results have suggested that the nucleation of specific intermediate oligomers is an essential aspect of canonical mature core assembly. It is also notable that self-assembly is predicted to proceed even in the absence of viral RNA, which is consistent with recent experiments that have demonstrated mature capsid formation when co-assembled with inositol phosphate, a small-molecule co-factor.^39^ The conditions used in previous simulations implicitly include the putative effects of these co-factors, and the simulations presented hereafter similarly assume that RNA is not essential for core formation.

**Figure 1.**
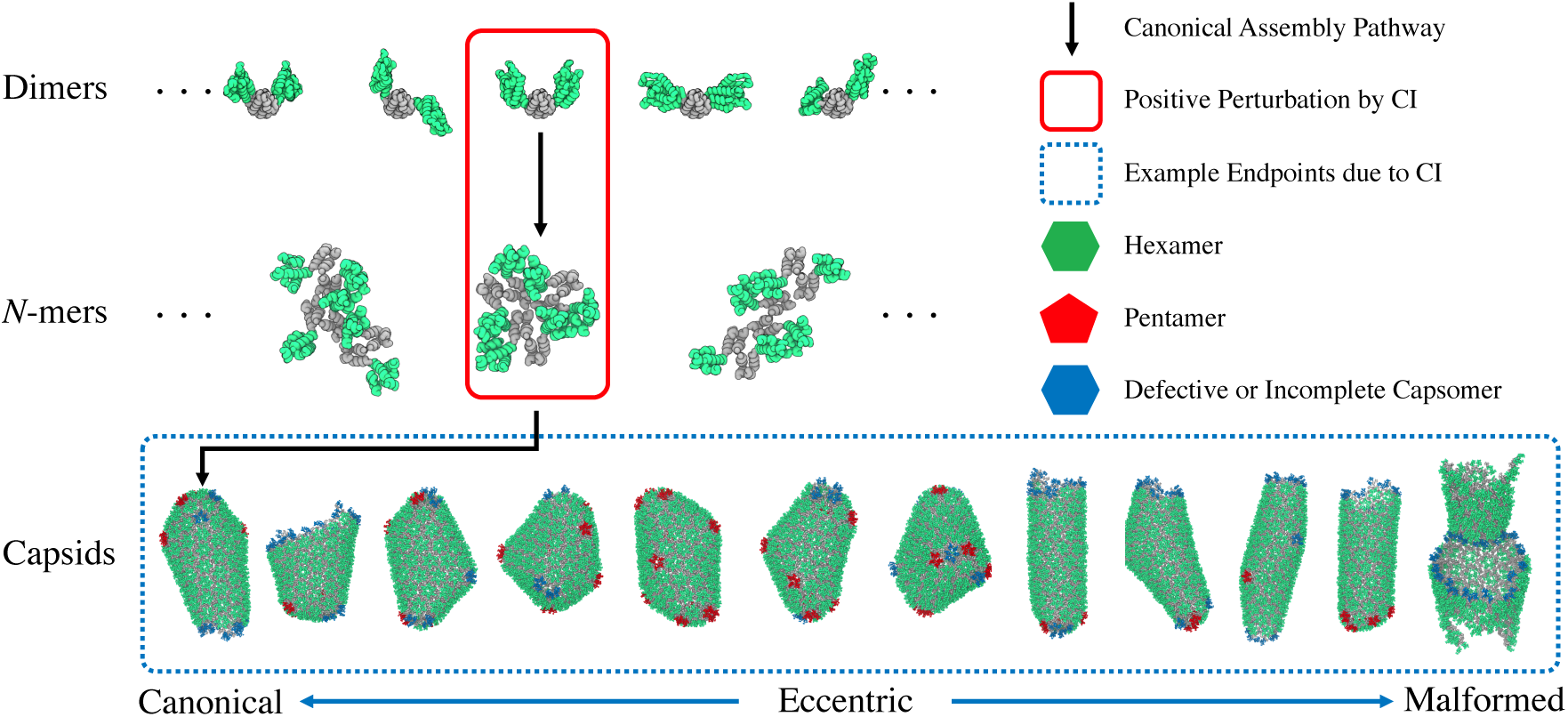
Schematic of the hierarchical process that is central to HIV-1 mature capsid assembly. Within each oligomeric state of *N* monomers (*N*-mer), a variety of configurations are possible. The canonical assembly pathway relies on constant self-correction across *N*-mer states, which is contingent on a dynamic and broad population of small *N*-mer intermediates. In this work, we effectively introduce the presence of capsid inhibitor (CI) drugs using a small but fixed population of trimers of dimers, a 6-mer with up to three bound CIs (one at each dimer-dimer interface), thereby perturbing the natural dynamics of the assembly process. Snapshots of the final structures from 12 CG-MD simulations are depicted, from which a majority population of eccentric or malformed capsids can be seen. Here, eccentric (canonical) end-points refer to structures with regions of densely-accumulated pentamers and defective hexamers (broadly distributed pentamers) while malformed assemblies are non-enclosed and semi-amorphous structures.

To consider the impact of CIs on the CA assembly process, we note that solved crystallographic structures show evidence that PF74 and GS-CA1 preferentially bind to an inter-CA pocket between the respective NTD and CTD domains of two adjacent CA monomers.^23, 25, 28^ Importantly, these pockets only appear when CA oligomerizes, such as in TODs, and appear to mediate inter-CA interactions, thereby stabilizing multimeric CA.^31^ We therefore hypothesize that CIs stabilize the interface between CA domains, increasing the population and/or lifetime of intermediate oligomers. A natural question to investigate is how these perturbations affect the overall CA assembly process.

To simulate the influence of CIs, we use the “Ultra-CG” (UCG) model^11^ that was previously developed to investigate mature capsid assembly. Briefly, the UCG model consists of a Cα resolution representation of CA helices with an elastic network model^40^ (ENM) used to conserve the tertiary structure of the protein. Select inter-CA contacts that are exposed in the “mature” configuration are identified from X-ray structures^18, 41^ and projected as “virtual” CG sites with two possible states: (1) a non-interacting “inactive” state and (2) an interacting (through an attractive Gaussian interaction) “active” state. Throughout the simulation, CA dimers periodically switch between active ([CA^+^]) and inactive ([CA^-^]) states and a constant ratio between the two is maintained. Here, we include the effect of CI binding to CA by introducing a fixed population of [CA^+^] that have preassembled into TODs ([CA^+^]_CI_), as highlighted by the red box in Fig. 1. We performed CG MD simulations under conditions that previously resulted in self-regulated assembly^11^ ([CA^+^]/[CA^-^] = 0.11, [CA] = 4 mM, inert crowder density at 200 mg/ml, UCG state switching interval at 5×10^5^ timesteps) with [CA^+^]_CI_ = 0.025, 0.050, 0.075, 0.100, 0.200 and 0.500 mM and 2 replicates for each condition resulting in a total of 12 independent trajectories; note that [CA^+^]_CI_ = 0.025 mM represents a single TOD out of a total of 616 CA dimers in a 80 nm length cubic box. Each simulation was run for 2×10^9^ timesteps (τ) with τ = 10 fs in CG time (which is not to be confused with actual time). Further details can be found in Methods and in Ref. 11.

The bottom row in Fig. 1 depicts the outcomes of each of the 12 CG simulations. Here, we qualitatively define three classes of core morphologies for the purposes of the following discussion: (1) canonical, (2) eccentric, and (3) malformed. Malformed capsids are those with non-contiguous, semi-amorphous lattices, such as when multiple aggregates of CA merge. Canonical and eccentric capsids refer to contiguous lattices that are distinguished by their curvatures, and relatedly, their pentamer distributions; in the former case, pentamers are dispersed such that all pentamers are separated by (at minimum) a hexamer, while the latter case contains adjacent pentamers in pentamer-dense regions, which also denote regions of high curvature; the need for this distinction will become evident during the discussion below. From our CG simulations, we observe 1 canonical, 10 eccentric, and 1 malformed capsid structure. We note that we do not observe any explicit trends between [CA^+^]_CI_ and the resultant core morphologies, as summarized in Table 1, which is likely due to the stochastic nature of the process; additional independent sampling at each [CA^+^]_CI_ would be required to confirm a potential trend with respect to [CA^+^]_CI_, but is currently computationally cost prohibitive and outside the scope of this work. Nonetheless, our simulations suggest that the majority of cores that form due to CIs are highly pleomorphic with many cores exhibiting large degrees of curvature. This is a notable result when compared to the expected behavior of WT viruses. Previous cryo-electron microscopy (cryo-EM) surveys of mature WT virions have revealed a variety of capsid phenotypes, including conical capsids, tubular capsids, incomplete capsids and dual capsids, although the majority (around 60 to 90%) appears to be conical.^42–44^ Our simulations suggest that the presence of CIs shifts this distribution toward the formation of non-conical cores, which is consistent with cryo-EM images of virions treated by GS-CA1.^25^ The distribution of end-point morphologies show that even slight positive perturbation to TOD populations has profound effects on CA assembly that favor alternative pathways. We note that this view is conceptually consistent with recent reports of pathway complexity observed in supramolecular polymers, which emerges when assembly kinetics cannot be described by simple nucleation-and-elongation models.^45–47^

**Table 1.** Summary of end-point capsid morphologies for each of the 12 listed systems. The numerical label in brackets refers to its associated snapshot in Fig. 1 with left-most = 1 and right-most = 12.

| $[\text{CA}^+]_{\text{CI}} \text{ (mM)}$ | Set 1 | Set 2 |
| --- | --- | --- |
| 0.025 | Eccentric, Closed [3] | Eccentric, Closed [6] |
| 0.050 | Canonical, Open [1] | Malformed [12] |
| 0.075 | Eccentric, Open [8] | Eccentric, Closed [7] |
| 0.100 | Eccentric, Closed [5] | Eccentric, Open [10] |
| 0.200 | Eccentric, Open [9] | Eccentric, Open [11] |
| 0.500 | Eccentric, Closed [4] | Eccentric, Open [2] |

To characterize the lattice morphology, we first recognize that the CA lattice can be represented as a connected graph. One may conceive of each CA monomer as a vertex (*v*) on a graph (*G*) with an edge (*e*) between vertices signifying their proximal nature. Within the assembled CA lattice, *e* represents dimeric CA as well as adjacent intra-capsomer CA; in other words, each *v* is ideally connected to three neighboring *v* within *G*, which strictly represents the largest assembled CA lattice. Throughout the simulation trajectories, we use a distance criterion to construct this graph representation:

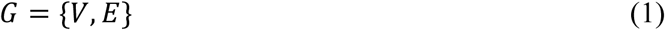

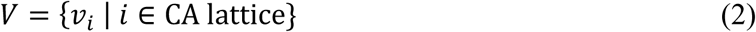

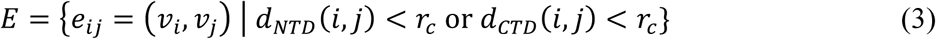

with *V* and *E* denoting the set of *v* and *e* in *G, i* and *j* denoting the index of each CA monomer, *d*_*NTD*_ (*d*_*CTD*_) denoting the measured distance between residue V36 (E180) in *i* and *j*, and *r*_*c*_ denoting a cutoff distance of 2.5 nm. We then consider a property of graphs known as eccentricity (*ecc*), which effectively measures how “far” a given vertex is from its furthest vertex. The *ecc* of each *v* is formally defined as:

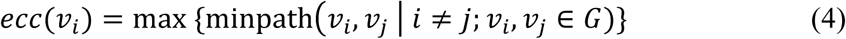

which quantifies the maximum path distance amongst the set of minimum paths between *v*_*i*_ and every other *v* in *V*. For our purposes, we consider the normalized *ecc* (λi = *ecc*(*v*_*i*_) / max{*ecc*(*v*)}) and statistical measures on its distribution (*P*(λ)) as a metric to classify CA lattice morphologies.

The utility of *P*(λ) can be shown with four representative CA lattices (extracted from a single trajectory) that are analyzed in Fig. 2. Fig. 2(a) depicts an early stage of CA nucleation in which the lattice appears largely two-dimensional and isotropic around its six-fold (C_6_) axis. The resultant *P*(λ) exhibits a normal distribution with the smallest (largest) λ_i_ associated with the interior (edges) of the lattice. In Fig. 2(b), the lattice has grown with some edges appearing “jagged” (a nascent dendrite, in some sense), which is indicative of anisotropic nucleation. The λ_i_ associated with *v* that participate in these jagged edges is large (e.g., λ_i_ > 0.8) in comparison to interior *v*; in fact, as these edges become more dendritic, we expect large λ_i_ to be increasingly favored in *P*(λ). Here, we argue that anisotropic growth at the edges of the CA lattice are relevant since the branching junction between dendrites may nucleate into pentamers, presumably to anneal the lattice and to minimize potential strain due to curvature. In Fig. 2(c), the lattice has fully wrapped into an enclosed 2D lattice, such that all *v* can be considered interior *v*. Now, *P*(λ) favors a higher population of small λ_i_ (e.g., λ_i_ < 0.8) with large λ_i_ associated with oblong areas of the capsid; note that pristine cleavage of the capsid, such that the edges are largely composed of hexamers, recovers a *P*(λ) with a normal distribution, as seen in Fig. 2(d). This analysis suggests that we may use the skew of *P*(λ) (i.e., γ(λ)) to qualitatively classify CA lattice growth; γ(λ) ≈ 0 signifies a lattice with edges that are isotropically distant from the lattice center while γ(λ) < 0 (γ(λ) > 0) represents a lattice with anisotropic edges (with contiguous and uniform oblong character, such as in complete enclosures).

**Figure 2.**
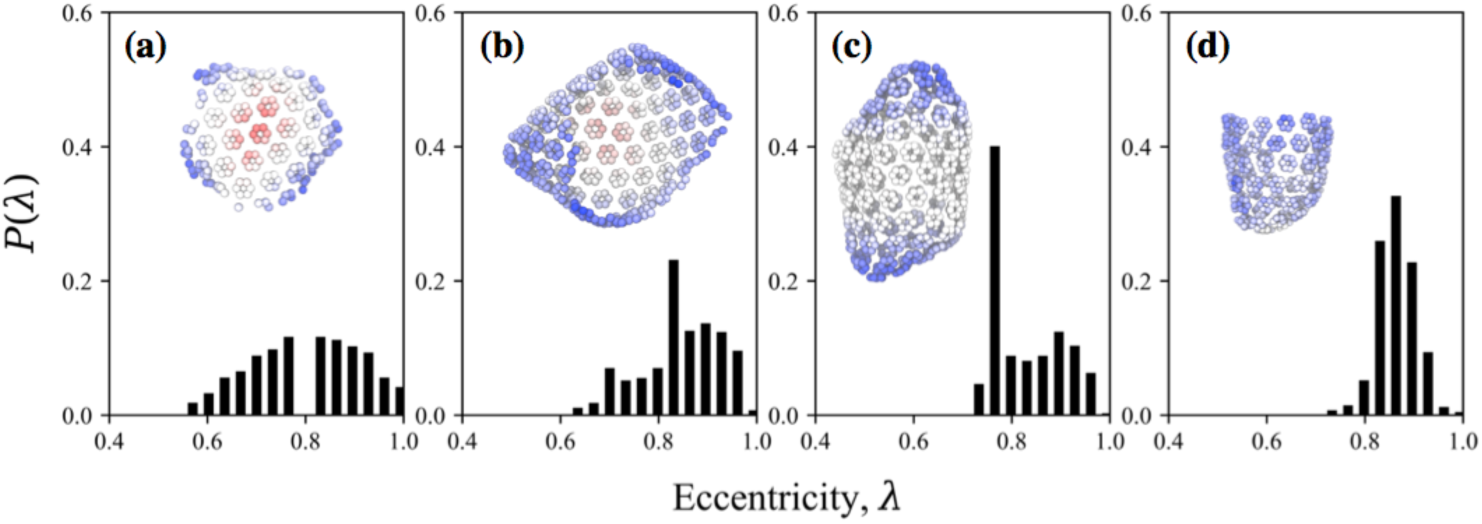
The topology of the assembled CA lattice is assessed by the eccentricity (λ) of each vertex (i.e., each CA monomer) in a graph representation of the lattice. We show the distribution of λ (*P*(λ)) at select points during one trajectory of [CA^+^]_CI_ = 0.100 mM for demonstrative purposes: at (a) 1×10^8^, (b) 7×10^8^, and (c) 14×10^8^ MD timesteps. The same analysis is performed in (d) using half of the lattice from (c), i.e., after cleavage by a plane perpendicular to the long axis of the capsid. Representative lattices are depicted as insets in (a-d) with each monomer shown as a sphere colored by its λ from red (λ=0.5) to blue (λ=1.0). The topology of the lattice can be qualitatively characterized by the skew of the distribution of λ (γ(λ)) in which (a,d) zero skew is indicative of an open lattice with isotropic edges, (b) negative skew is indicative of an open anisotropic lattice, and (c) positive skew is indicative of a closed, non-spherical lattice.

On the basis of the aforementioned metric, we now analyze time-series profiles from a subset of the CG trajectories, which were chosen for clarity, in Fig. 3 and compare them to that of the non-CI case from Ref. 11. We find that in all cases with [CA^+^]_CI_ > 0, the assembly rate for CA hexamers (Fig. 3(a)) and pentamers (Fig. 3(b)) is accelerated compared to the [CA^+^]_CI_ = 0 case. The observed increase in multimerization rate is consistent with experimental multimerization assays for PF74,^23, 31^ and our simulations suggest that CI-bound populations of CA (even at stoichiometric ratios as low as 1 mol%) can have a profound impact on multimerization rates. Interestingly, the onset of pentamer incorporation is also notably sooner in the CI-bound cases (around 50–150×10^6^ τ) compared to the CI-absent case (around 350×10^6^τ). In all cases, γ(λ), as seen in Fig. 3(c), is initially close to zero but shifts toward negative values directly preceding the onset of pentamer incorporation; recall that negative γ(λ) is indicative of non-uniform protrusions along the lattice edges. The resultant pentamer incorporation typically seems to anneal the lattice such that γ(λ) increases toward 0 or positive values and the rate of pentamer incorporation is temporarily suppressed. However, as the edges of the lattice continue to nucleate CA, anisotropic character may re-emerge and promote the additional incorporation of pentamers. The edges of the lattice may ultimately anneal to form an enclosed CA lattice (with γ(λ) > 0). Here, CA within incomplete capsomers, which are likely to be in areas of high curvature, continue to relax until pentamers are formed, e.g., as evident by the time-series profiles in which pentamer counts increase while hexamer growth is stalled. Molecular snapshots from a CI-bound case that exhibits all of these steps are shown in Fig. 3(d).

**Figure 3.**
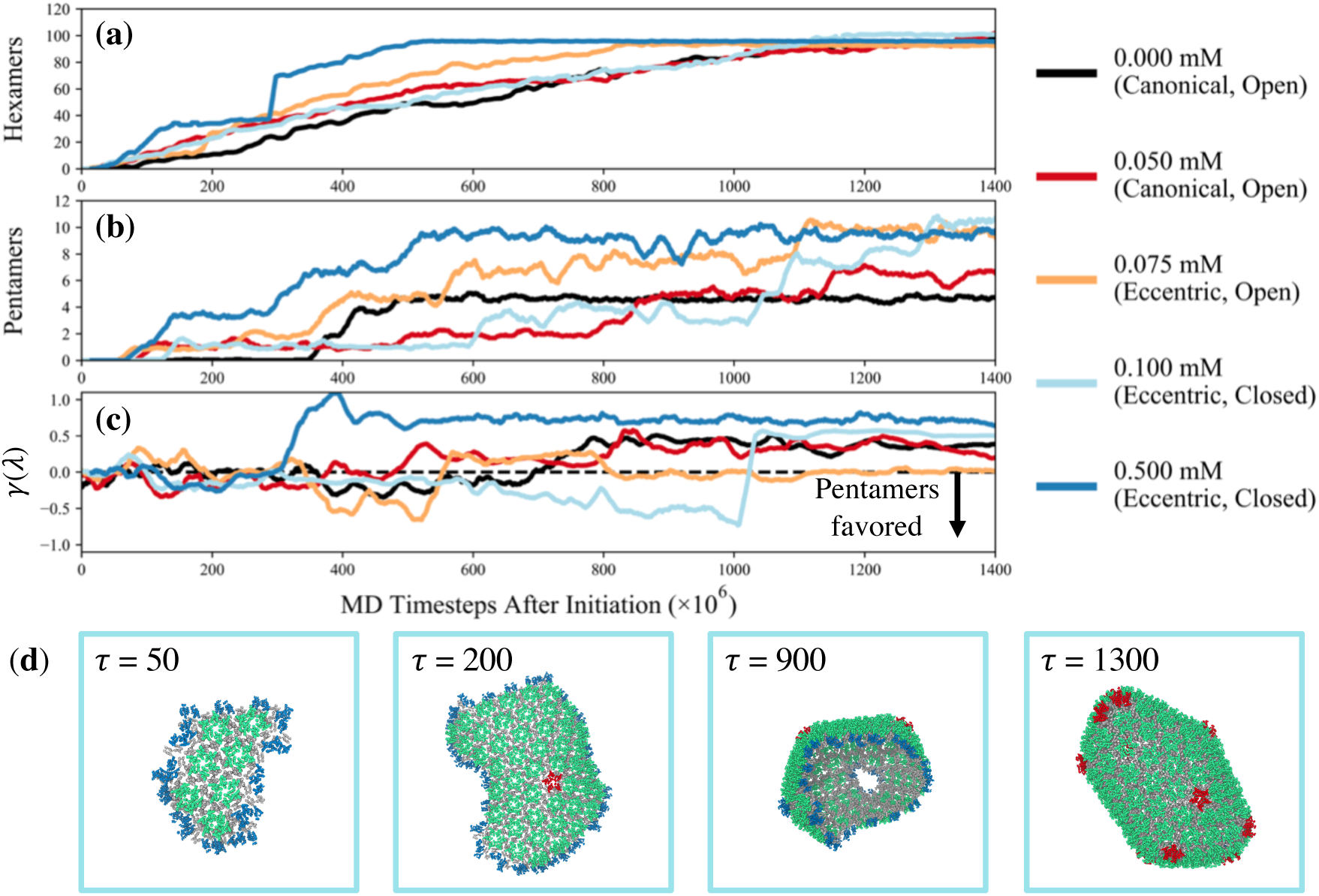
Assembly time-series plots that depict (a) the number of assembled hexamers, (b) the number of assembled pentamers, and (c) the skew (γ(λ)) of the distribution of eccentricities (λ) throughout the assembled lattice as a function of CG MD timestep (shifted with respect to the onset of lattice growth) for each system with the listed concentration of capsid inhibitors (CIs). We find accelerated assembly, especially with respect to pentamers, in the CI-present simulations, which appear to be commensurate with negative γ(λ), i.e., a descriptor that indicates anisotropic edge growth in the protein lattice. In (d), molecular snapshots of the protein lattice for the 0.100 mM case (cyan line in (a)-(c)) at the listed MD timestep (τ [×10^6^]) are shown; green, red, and blue NTD domains in capsomers indicate hexamers, pentamers, and incomplete capsomers, respectively.

We interpret these collective results by proposing that CI-bound TODs (i.e., the fixed TODs in our simulations) accelerate CA assembly by increasing the number of accessible assembly pathways. First, let us consider that TODs, while important building blocks, still require free CA dimers (and potentially, other small oligomers) to continue assembly of the CA lattice.^11^ Each CA dimer favors two binding interfaces that should be satisfied for successful association, as seen in Fig. 4(a). Hence, continued CA assembly requires the near-simultaneous fastening of two CA dimers at adjacent edges of the lattice, which subsequently create new edges for future second-order CA association events. The increased population of TODs, on the other hand, enables CA lattice growth through the near-simultaneous binding of a TOD and CA dimer. In this alternative assembly pathway, new binding edges also appear. Yet, it is instead likely that subsequent assembly steps only require a single CA dimer, as seen in Fig. 4(a), which therefore accelerates CA assembly.

**Figure 4.**
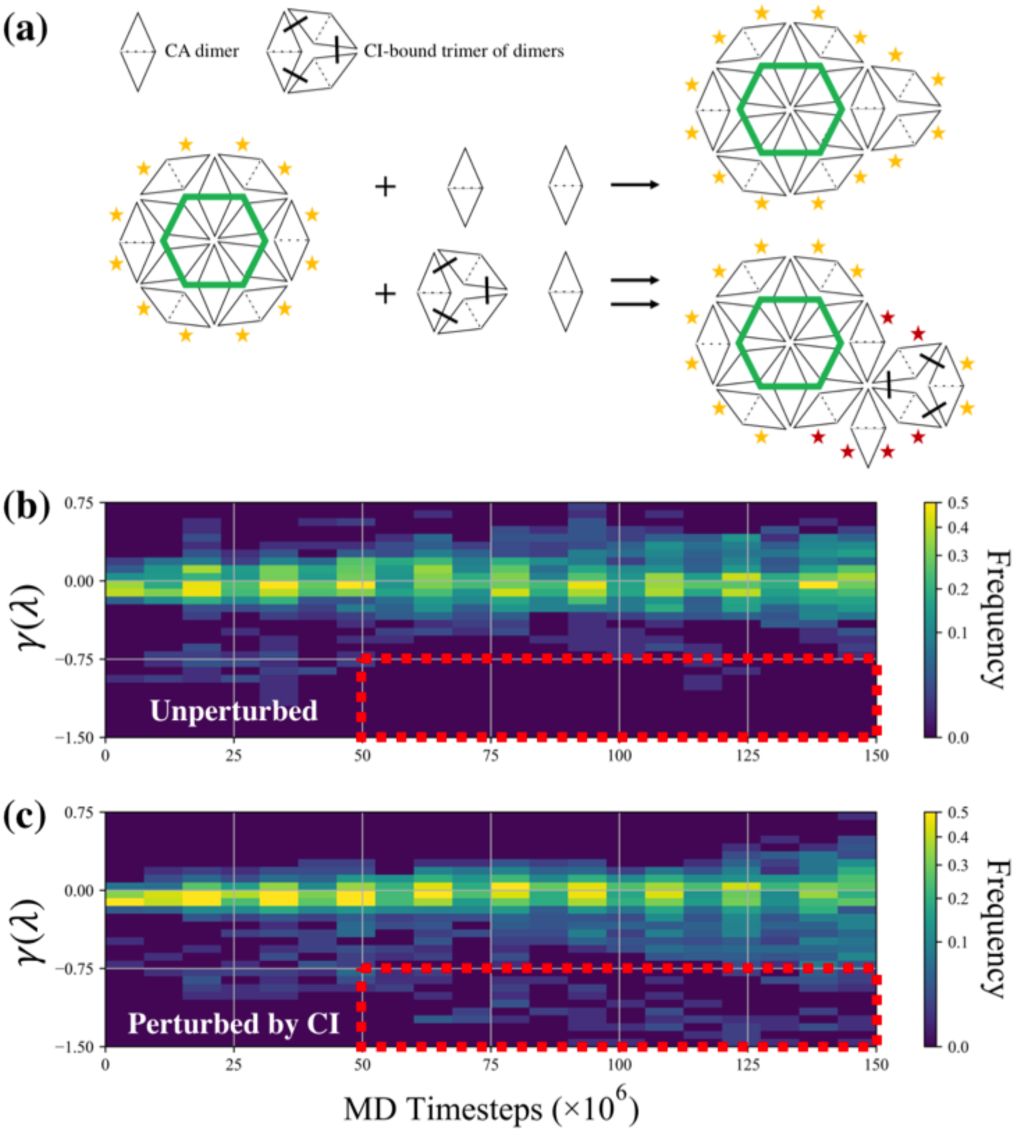
(a) Schematic of the canonical assembly pathway (top) through the dual association of CA dimers and accelerated assembly pathway (bottom) through the association of a trimer of dimer (TOD) that is stabilized by capsid inhibitors (CIs) and a CA dimer. The stars represent potential binding interfaces; yellow stars indicate the need for an adjacent CA dimer to associate nearly simultaneously, while red stars indicate sites that are energetically satisfied by a single CA dimer. The topological character of an assembling lattice over time is quantified by the skew of its eccentricities (γ(λ)) with distributions calculated from a swarm of 50 trajectories in the (b) absence and (c) presence of CIs. Here, γ(λ) ≈ 0 is indicative of an open isotropic lattice, γ(λ) < 0 is indicative of an open anisotropic lattice, and γ(λ) > 0 is indicative of a uniform and oblong lattice (such as in closed mature capsids). These results suggest the preference of isotropic (emergence of anisotropic) lattice growth during canonical (CI-perturbed) assembly, which is highlighted in the region bound by the dashed red box. Each trajectory was discretized into 20 equal bins before histograms were computed (*n* = 375 per histogram).

To demonstrate the emergence of additional assembly paths, we perform 50 short independent simulations (over 150×10^6^ τ) of CA assembly with [CA^+^]_CI_ = 0.000 and 0.100 mM starting from a pre-assembled lattice with 37 CA hexamers (i.e., the equivalent of four concentric “layers” of hexamers). We depict the progression of γ(λ) in Fig. 4(b-c) and compare the two cases. In the [CA^+^]_CI_ = 0.000 mM case (Fig. 4(b)), we find that the majority of trajectories maintain γ(λ) around 0.0. However, in the [CA^+^]_CI_ = 0.100 mM case (Fig. 4(c)), a fraction of the trajectories instead adopt γ(λ) < 0, which is commensurate with increasingly anisotropic lattice growth, with this fraction increasing as time progresses. This observation is most evident after 50×10^6^ τ has elapsed; here, states associated with γ(λ) < –0.75 are sampled by CI-bound trajectories while absent from apo trajectories (see red box in Fig. 4(b)). We therefore suggest that the presence of CI-bound TODs enables an alternative assembly pathway through enhanced CA association kinetics and increasingly anisotropic lattice growth.

One consequence of the accelerated CA assembly, and relatedly, the rapid pentamer incorporation, induced by CIs is that the resultant mature cores may fail to enclose viral RNA for two reasons. First, the enhanced CA assembly kinetics may preclude the need for viral RNA and nucleocapsid (NC), i.e., the ribonucleoprotein (RNP) complex, to serve as templates for assembly; some experiments have suggested that the RNP complex nucleates mature CA assembly,^42^ which is further mediated by integrase,^43, 48^ although this behavior does not appear to be universal as evident by mature capsid formation even in the absence of an enclosed RNP complex.^43–44, 49^ Second, CI-perturbed cores are likely to adopt highly curved morphologies with small internal volumes that may impede the encapsulation of the RNP complex. In our simulations, cores that underwent full enclosure were observed to contain on the order of 100 hexamers (see Fig. 3(a)). For comparison, previous accounts from cryo-EM data suggest canonical capsid cores contain on the order of 200 hexamers. Based on an effective reduction of surface area by a factor of two, we approximate an associated reduction of internal volume by up to 65% (for simple spherical geometries). However, we note that RNA condensation, which is likely dependent on salt and viral protein content, may allow the RNP complex to occupy these reduced volumes; note that such condensation is suggested by the localization of RNP-associated densities in cryo-EM images within the broad region of conical mature capsids.^42, 49^ Nonetheless, the proposed phenotype is consistent with previous cryo-EM experiments that have observed RNP-associated density outside that of protein cores, especially those treated with inhibitors.^43–44^ Hence, our simulations suggest that one mechanism of action of CIs is to increase the population of eccentric cores that may fail to enclose the RNP complex.

In the event that CI-bound mature cores successfully condense the RNP complex, we also investigated their viral uncoating behavior. During the post-entry stages of viral infection, capsid disassembly is highly regulated.^6^ Although the process is not completely understood, it has been suggested that certain biological triggers, such as the binding of host cell factors^50–52^ or the initiation of reverse transcription,^53^ act as signals to initiate disassembly. Hence, perturbations to canonical uncoating behavior may result in reduced infectivity. We simulate putative post-entry events following the procedures established in Ref. 11 in which we deplete the concentration of available [CA^+^] (= 0 mM) and integrate dynamics over 8×10^8^ τ. In Fig. 5, we compare time-series profiles that depict the fraction of the mature core that persists; here, we select and compare four closed end-point morphologies from our simulations described above. Interestingly, three out of four of the selected eccentric cores undergo a step-wise disassembly process, as described by the processive periods of plateaus and decrements seen in Fig. 5(a). The molecular snapshots depicted in Fig. 5(b) of an eccentric core during disassembly provide insight into the origins of this behavior. During the plateau periods, pentamers switch between pentameric states (red in Fig. 5(b)) and that of an incomplete hexamer (blue in Fig. 5(b)). In this latter state, it is more likely for a CA dimer to dissociate from the lattice, which has been suggested as the rate-limiting step for viral uncoating. ^54^ The resultant defect is then the site of spontaneous dissociation of CA. In comparison, we find that one of the cores remains enclosed, which is evident from the flat profile in Fig. 5(a). This suggests that within our simulated time-scales, some eccentric cores are inherently stable. In this case, we find that the regions with large pentamer density are restricted to two adjacent pentamers at most, and therefore speculate that an increased local density of pentamers (of three or more) is required for spontaneous uncoating. These results are consistent with fluorescence experiments that have suggested both WT and PF74-treated viruses contain small populations of cores that fail to open, while the remaining PF74-treated viral cores have a greater propensity to open spontaneously.^54^ Another mechanism of action for CIs may therefore be the production of eccentric cores that exhibit rapid CA disassembly due to inherent instabilities in high curvature regions with large pentamer density. Hence, despite the successful assembly of mature cores in the presence of CIs, accelerated capsid disassembly during the early stages of infection may inhibit viral infectivity.

**Figure 5.**
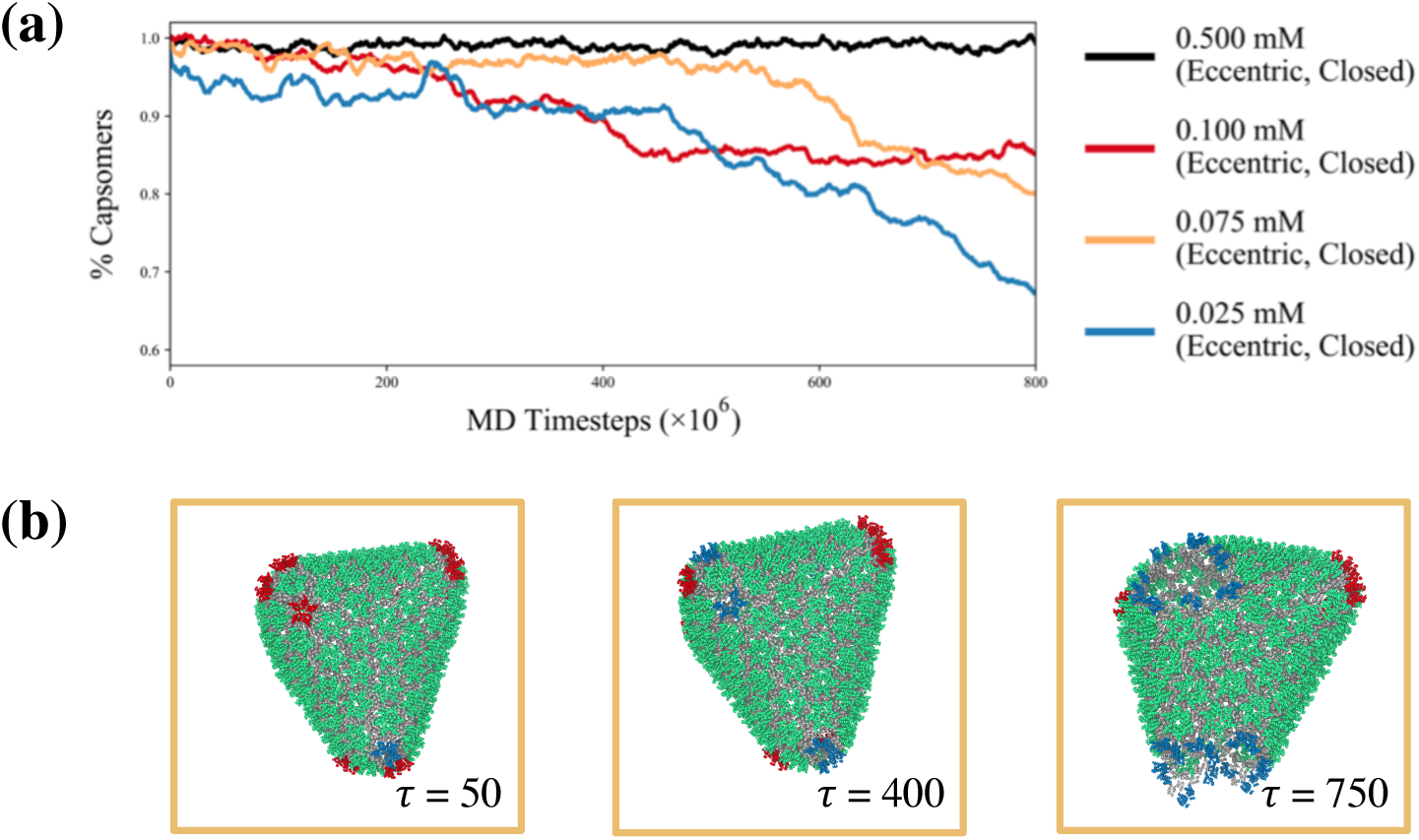
(a) Disassembly time-series plot that depicts the fraction of remaining hexamers and pentamers (with respect to the initial size of the mature capsid core) for the four listed systems as a function of MD timestep. Three of the eccentric cases exhibit spontaneous disassembly, which appear to be initiated at sites with large pentamer density. (b) Molecular snapshots of the protein lattice for the 0.075 mM case (orange line in (a)) at the listed MD timestep (τ [×10^6^]) are shown; green, red, and blue capsomers indicate hexamers, pentamers, and incomplete capsomers, respectively. This suggests that capsid inhibitors can promote the formation of ‘leaky’ mature cores that spontaneously open at regions of high curvature (or high pentamer density), and represents one failure mechanism that suppresses infectivity.

On a final note, our simulations demonstrate that a CG model can aid interpretation of the anomalously broad activity of CIs observed in experiments. In the case of PF74, the binding of CIs to pre-established binding pockets stabilizes CA multimers in tubular structures and prevents disassembly.^28, 30–31^ Our simulations incorporate this stabilization effect with the use of TODs that are fixed during dynamics. We only consider a low stoichiometric ratio of CI-bound CA given that experimental EC_50_ estimates tend to be in the range of µM concentrations or below.^23–25^ Our results show that perturbing TOD distributions in this way is sufficient to induce alternative assembly pathways that yield diverse capsid morphologies, including eccentric cores that were previously seen as a minority population in WT viruses.^42–44^ The inherent diversity of these capsids results in multiple mechanisms of action. Furthermore, these mechanisms appear to be fundamentally different from that of inhibitors that target monomeric CA, which tend to affect either early or late (but not both) stages of the viral lifecycle. For example, CAP-1 and CAI bind to CA monomers and prevent proper CA association at key interfaces (e.g., the CTD dimeric interface) during late-stage assembly.^55–56^ For these types of CIs, higher drug concentrations (e.g. µM or above) may have been required to ensure that the population of apo CA monomers is sufficiently repressed to prevent complete assembly. Hence, CIs that target the highly regulated dynamical processes observed throughout the viral lifecycle by means of CA oligomer subpopulations may have two primary benefits: (1) broader spectrum inhibitory effects and (2) reduced stoichiometric requirements that facilitate drug delivery challenges.

## Conclusions

In summary, we use CG molecular dynamics simulations and elucidate the mechanisms underlying the broad-spectrum HIV-1 inhibitory effects of capsid inhibitors (CIs), such as PF74 and GS-CA1, by virtue of small perturbations to the hierarchical self-assembly of viral capsid (CA) proteins. We find that the primary action of CIs is to stimulate CA association and propagate anisotropic assembly pathways, which results from the stabilization of a subpopulation of trimer of dimers (TODs); this behavior is consistent in all of our CG simulations, even at the lowest accessible drug to CA stoichiometries of less than 1 mol%. Consequently, pentameric defects appear to be rapidly incorporated, thereby increasing the expression of eccentric cores, that is, mature cores with inherently large curvature and pleomorphism. Capsids of this phenotype may unsuccessfully enclose the viral nucleic acid and nucleocapsid protein (RNP) complex, which is one possible failure mode for infection. However, in the event that the RNP complex is encapsulated, a secondary failure mode during post-entry events is observed from our capsid disassembly simulations. Here, regions of the capsid that are dense with pentameric defects appear to be inherently unstable and initialize CA dissociation; eccentric cores that form due to CIs may therefore spontaneously disassemble after entry into the cytoplasm. Taken together, our findings reveal that CIs affect processes during both viral maturation and uncoating. These results also suggest that the identification of potential binding pockets that appear upon CA oligomerization, especially those with small degrees of genetic polymorphism, and the design of small molecules that specifically target these sites – even at low stoichiometric loadings – will facilitate the design of HIV-1 (and other viral) inhibitors with similarly broad mechanisms of action.

## Methods

### Coarse-grained modeling and simulation

As mentioned above, the CG models used in this work are described in Ref. ^11^. Our simulations contained CA, pre-assembled TODs and inert crowding agents, as described in the main text, which were initially dispersed randomly within the periodic simulation box. Configurations were allowed to equilibrate over 5×10^6^ τ without any attractive interactions. Production runs were then performed as described in the main text, while [CA^+^]/[CA^-^] UCG state switching was attempted every 5×10^5^ τ. All simulations were performed in the constant *NVT* ensemble at 300 K using a Langevin thermostat^57^ with a damping period of 100 ps and τ = 10 fs. All simulations were run using an in-house MD engine called UCG-MD, which is optimized to run implicit-solvent UCG simulations with dynamic load-balancing and runtime algorithms.^58^ Neighbor lists up to 4 nm with an additional 1 nm skin depth were used, while load balancing was attempted every 2×10^4^ τ. Molecular snapshots were saved every 1×10^6^ τ for analysis.

### Data analysis

Graph analysis was performed using the python package NetworkX 2.1 (http://networkx.github.io/). A graph was constructed for each molecular snapshot following the criteria described in the main text. Hexamers and pentamers were identified by vertices (i.e., CA monomers) that participated in a closed cycle of length six and five, respectively. The remaining nodes were identified as components of incomplete capsomers. The λ of each vertex was computed using the *eccentricity* function.

## Acknowledgements

Support from the Ruth L. Kirschstein National Research Service Award Postdoctoral Fellowship is gratefully acknowledged by AJP (grant F32-GM125218). This work was also supported by the National Institute of General Medical Sciences of the National Institutes of Health (grants R01-GM128507 and P50-GM082545 for GAV, JMAG, and AY).

## References

1. Arts, E. J.; Hazuda, D. J., HIV-1 Antiretroviral Drug Therapy. Csh Perspect Med 2012, 2 (4).

2. Cihlar, T.; Fordyce, M., Current status and prospects of HIV treatment. Curr Opin Virol 2016, 18, 50–56.

3. Laskey, S. B.; Siliciano, R. F., A mechanistic theory to explain the efficacy of antiretroviral therapy. Nature Reviews Microbiology 2014, 12 (11), 772–780.

4. Sundquist, W. I.; Kräusslich, H. G., HIV-1 assembly, budding, and maturation. Csh Perspect Med 2012, 2, a006924.

5. Freed, E. O., HIV-1 assembly, release and maturation. Nature Reviews Microbiology 2015, 13, 484–496.

6. Ambrose, Z.; Aiken, C., HIV-1 uncoating: connection to nuclear entry and regulation by host proteins. Virology 2014, 454-455, 371–379.

7. Adamson, C. S.; Sakalian, M.; Salzwedel, K.; Freed, E. O., Polymorphisms in Gag spacer peptide 1 confer varying levels of resistance to the HIV-1maturation inhibitor bevirimat. Retrovirology 2010, 7 (1), 36.

8. Bell, N. M.; Lever, A. M. L., HIV Gag polyprotein: Processing and early viral particle assembly. Trends Microbiol. 2013, 21, 136–144.

9. Freed, E. O., HIV-1 Gag proteins: Diverse functions in the virus life cycle. Virology 1998, 251, 1–15.

10. Grime, J. M. A.; Voth, G. A., Early stages of the HIV-1 capsid protein lattice formation. Biophys. J. 2012, 103, 1774–1783.

11. Grime, J. M. A.; Dama, J. F.; Ganser-Pornillos, B. K.; Woodward, C. L.; Jensen, G. J.; Yaeger, M. J.; Voth, G. A., Coarse--grained simulation reveals key features of HIV--1 capsid self-assembly. Nat Commun 2016, 7, 11568.

12. Pak, A. J.; Grime, J. M. A.; Sengupta, P.; Chen, A. K.; Durumeric, A. E. P.; Srivastava, A.; Yeager, M.; Briggs, J. A. G.; Lippincott-Schwartz, J.; Voth, G. A., Immature HIV-1 lattice assembly dynamics are regulated by scaffolding from nucleic acid and the plasma membrane. P Natl Acad Sci USA 2017, 114 (47), E10056–E10065.

13. Bharat, T. A. M.; Castillo Menendez, L. R.; Hagen, W. J. H.; Lux, V.; Igonet, S.; Schorb, M.; Schur, F. K. M.; Kräusslich, H.-G.; Briggs, J. A. G., Cryo-electron microscopy of tubular arrays of HIV-1 Gag resolves structures essential for immature virus assembly. Proceedings of the National Academy of Sciences 2014, 111, 8233–8.

14. Schur, F. K. M.; Hagen, W. J. H.; Rumlová, M.; Ruml, T.; Müller, B.; Kräusslich, H.-G.; Briggs, J. A. G., Structure of the immature HIV-1 capsid in intact virus particles at 8.8 Å resolution. Nature 2015, 517, 505–508.

15. Schur, F. K. M.; Obr, M.; Hagen, W. J. H.; Wan, W.; Jakobi, A. J.; Kirkpatrick, J. M.; Sachse, C.; Kräusslich, H.-g.; Briggs, J. A. G., An atomic model of HIV-1 capsid-SP1 reveals structures regulating assembly and maturation. Science 2016, 353, 506–508.

16. Wagner, J. M.; Zadrozny, K. K.; Chrustowicz, J.; Purdy, M. D.; Yeager, M.; Ganser-Pornillos, B. K.; Pornillos, O., Crystal structure of an HIV assembly and maturation switch. Elife 2016, 5, e17063.

17. Ganser, B. K.; Li, S.; Klishko, V. Y.; Finch, J. T.; Sundquist, W. I., Assembly and analysis of conical models for the HIV-1 core. Science 1999, 283, 80–83.

18. Pornillos, O.; Ganser-Pornillos, B. K.; Yeager, M., Atomic-level modelling of the HIV capsid. Nature 2011, 469, 424–7.

19. Li, G.; Verheyen, J.; Rhee, S.-Y.; Voet, A.; Vandamme, A.-M.; Theys, K., Functional conservation of HIV-1 Gag: implications for rational drug design. Retrovirology 2013, 10 (1), 126.

20. Rihn, S. J.; Wilson, S. J.; Loman, N. J.; Alim, M.; Bakker, S. E.; Bhella, D.; Gifford, R. J.; Rixon, F. J.; Bieniasz, P. D., Extreme Genetic Fragility of the HIV-1 Capsid. PLoS Path. 2013, 9 (6), e1003461.

21. Thenin-Houssier, S.; Valente, S. T., HIV-1 Capsid Inhibitors as Antiretroviral Agents. Curr. HIV Res. 2016, 14 (3), 270–282.

22. Carnes, S. K.; Sheehan, J. H.; Aiken, C., Inhibitors of the HIV-1 capsid, a target of opportunity. Curr Opin HIV Aids 2018, 13 (4), 359–365.

23. Blair, W. S.; Pickford, C.; Irving, S. L.; Brown, D. G.; Anderson, M.; Bazin, R.; Cao, J. A.; Ciaramella, G.; Isaacson, J.; Jackson, L.; Hunt, R.; Kjerrstrom, A.; Nieman, J. A.; Patick, A. K.; Perros, M.; Scott, A. D.; Whitby, K.; Wu, H.; Butler, S. L., HIV Capsid is a Tractable Target for Small Molecule Therapeutic Intervention. PLoS Path. 2010, 6 (12).

24. Li, F.; Goila-Gaur, R.; Salzwedel, K.; Kilgore, N. R.; Reddick, M.; Matallana, C.; Castillo, A.; Zoumplis, D.; Martin, D. E.; Orenstein, J. M.; Allaway, G. P.; Freed, E. O.; Wild, C. T., PA-457: A potent HIV inhibitor that disrupts core condensation by targeting a late step in Gag processing. Proceedings of the National Academy of Sciences 2003, 100 (23), 13555.

25. Tse, W., Discovery of novel potent HIV capsid inhibitors with long-acting potential. In Conference on Retroviruses and Opportunistic Infections, Seattle, Washington, 2017.

26. Shi, J.; Zhou, J.; Shah, V. B.; Aiken, C.; Whitby, K., Small-Molecule Inhibition of Human Immunodeficiency Virus Type 1 Infection by Virus Capsid Destabilization. J. Virol. 2011, 85 (1), 542.

27. Xu, H.; Franks, T.; Gibson, G.; Huber, K.; Rahm, N.; De Castillia, C. S.; Luban, J.; Aiken, C.; Watkins, S.; Sluis-Cremer, N.; Ambrose, Z., Evidence for biphasic uncoating during HIV-1 infection from a novel imaging assay. Retrovirology 2013, 10 (1), 70.

28. Bhattacharya, A.; Alam, S. L.; Fricke, T.; Zadrozny, K.; Sedzicki, J.; Taylor, A. B.; Demeler, B.; Pornillos, O.; Ganser-Pornillos, B. K.; Diaz-Griffero, F.; Ivanov, D. N.; Yeager, M., Structural basis of HIV-1 capsid recognition by PF74 and CPSF6. P Natl Acad Sci USA 2014, 111 (52), 18625–18630.

29. Hulme, A. E.; Kelley, Z.; Foley, D.; Hope, T. J., Complementary Assays Reveal a Low Level of CA Associated with Viral Complexes in the Nuclei of HIV-1-Infected Cells. J. Virol. 2015, 89 (10), 5350.

30. Fricke, T.; Brandariz-Nuñez, A.; Wang, X.; Smith, A. B.; Diaz-Griffero, F., Human Cytosolic Extracts Stabilize the HIV-1 Core. J. Virol. 2013, 87 (19), 10587.

31. Lad, L.; Clancy, S.; Koditek, D.; Wong, M. H.; Jin, D.; Niedziela-Majka, A.; Papalia, G. A.; Hung, M.; Yant, S.; Somoza, J. R.; Hu, E.; Chou, C.; Tse, W.; Halcomb, R.; Sakowicz, R.; Pagratis, N., Functional Label-Free Assays for Characterizing the in Vitro Mechanism of Action of Small Molecule Modulators of Capsid Assembly. Biochemistry-Us 2015, 54 (13), 2240–2248.

32. Price, A. J.; Jacques, D. A.; McEwan, W. A.; Fletcher, A. J.; Essig, S.; Chin, J. W.; Halambage, U. D.; Aiken, C.; James, L. C., Host Cofactors and Pharmacologic Ligands Share an Essential Interface in HIV-1 Capsid That Is Lost upon Disassembly. PLoS Path. 2014, 10 (10), e1004459.

33. Brass, A. L.; Dykxhoorn, D. M.; Benita, Y.; Yan, N.; Engelman, A.; Xavier, R. J.; Lieberman, J.; Elledge, S. J., Identification of Host Proteins Required for HIV Infection Through a Functional Genomic Screen. Science 2008, 319 (5865), 921.

34. König, R.; Zhou, Y.; Elleder, D.; Diamond, T. L.; Bonamy, G. M. C.; Irelan, J. T.; Chiang, C.-y.; Tu, B. P.; De Jesus, P. D.; Lilley, C. E.; Seidel, S.; Opaluch, A. M.; Caldwell, J. S.; Weitzman, M. D.; Kuhen, K. L.; Bandyopadhyay, S.; Ideker, T.; Orth, A. P.; Miraglia, L. J.; Bushman, F. D.; Young, J. A.; Chanda, S. K., Global Analysis of Host-Pathogen Interactions that Regulate Early-Stage HIV-1 Replication. Cell 2008, 135 (1), 49–60.

35. Deshmukh, L.; Ghirlando, R.; Clore, G. M., Conformation and dynamics of the Gag polyprotein of the human immunodeficiency virus 1 studied by NMR spectroscopy. Proceedings of the National Academy of Sciences 2015, 112, 3374–3379.

36. Deshmukh, L.; Schwieters, C. D.; Grishaev, A.; Ghirlando, R.; Baber, J. L.; Clore, G. M., Structure and dynamics of full-length HIV-1 capsid protein in solution. Journal of American Chemical Society 2013, 135, 16133–16147.

37. Qiao, X.; Jean, J.; Weber, J.; Zhu, F. Q.; Chen, B., Mechanism of polymorphism and curvature of HIV capsid assemblies probed by 3D simulations with a novel coarse grain model. Bba-Gen Subjects 2015, 1850 (11), 2353–2367.

38. Tsiang, M.; Niedziela-Majka, A.; Hung, M.; Jin, D. B.; Hu, E.; Yant, S.; Samuel, D.; Liu, X. H.; Sakowicz, R., A Trimer of Dimers Is the Basic Building Block for Human Immunodeficiency Virus-1 Capsid Assembly. Biochemistry-Us 2012, 51 (22), 4416–4428.

39. Dick, R. A.; Zadrozny, K. K.; Xu, C.; Schur, F. K. M.; Lyddon, T. D.; Ricana, C. L.; Wagner, J. M.; Perilla, J. R.; Ganser-Pornillos, B. K.; Johnson, M. C.; Pornillos, O.; Vogt, V. M., Inositol phosphates are assembly co-factors for HIV-1. Nature 2018, 560 (7719), 509–512.

40. Tirion, M. M., Large amplitude elastic motions in proteins from a single-parameter, atomic analysis. Phys. Rev. Lett. 1996, 77 (9), 1905–1908.

41. Pornillos, O.; Ganser-Pornillos, B. K.; Kelly, B. N.; Hua, Y.; Whitby, F. G.; Stout, C. D.; Sundquist, W. I.; Hill, C. P.; Yeager, M., X-Ray Structures of the Hexameric Building Block of the HIV Capsid. Cell 2009, 137 (7), 1282–1292.

42. Briggs, J. A. G.; Wilk, T.; Welker, R.; Kräusslich, H. G.; Fuller, S. D., Structural organization of authentic, mature HIV-1 virions and cores. The EMBO Journal 2003, 22 (7), 1707.

43. Fontana, J.; Jurado, K. A.; Cheng, N. Q.; Ly, N. L.; Fuchs, J. R.; Gorelick, R. J.; Engelman, A. N.; Steven, A. C., Distribution and Redistribution of HIV-1 Nucleocapsid Protein in Immature, Mature, and Integrase-Inhibited Virions: a Role for Integrase in Maturation. J. Virol. 2015, 89 (19), 9765–9780.

44. Wang, W. F.; Zhou, J.; Halambage, U. D.; Jurado, K. A.; Jamin, A. V.; Wang, Y. J.; Engelman, A. N.; Aiken, C., Inhibition of HIV-1 Maturation via Small-Molecule Targeting of the Amino-Terminal Domain in the Viral Capsid Protein. J. Virol. 2017, 91 (9).

45. van der Zwaag, D.; Pieters, P. A.; Korevaar, P. A.; Markvoort, A. J.; Spiering, A. J. H.; de Greef, T. F. A.; Meijer, E. W., Kinetic Analysis as a Tool to Distinguish Pathway Complexity in Molecular Assembly: An Unexpected Outcome of Structures in Competition. J. Am. Chem. Soc. 2015, 137 (39), 12677–12688.

46. Korevaar, P. A.; George, S. J.; Markvoort, A. J.; Smulders, M. M. J.; Hilbers, P. A. J.; Schenning, A. P. H. J.; De Greef, T. F. A.; Meijer, E. W., Pathway complexity in supramolecular polymerization. Nature 2012, 481, 492.

47. Haedler, A. T.; Meskers, S. C. J.; Zha, R. H.; Kivala, M.; Schmidt, H.-W.; Meijer, E. W., Pathway Complexity in the Enantioselective Self-Assembly of Functional Carbonyl-Bridged Triarylamine Trisamides. J. Am. Chem. Soc. 2016, 138 (33), 10539–10545.

48. Kessl, J. J.; Kutluay, S. B.; Townsend, D.; Rebensburg, S.; Slaughter, A.; Larue, R. C.; Shkriabai, N.; Bakouche, N.; Fuchs, J. R.; Bieniasz, P. D.; Kvaratskhelia, M., HIV-1 Integrase Binds the Viral RNA Genome and Is Essential during Virion Morphogenesis. Cell 2016, 166 (5), 1257-+.

49. Woodward, C. L.; Cheng, S. N.; Jensen, G. J., Electron Cryotomography Studies of Maturing HIV-1 Particles Reveal the Assembly Pathway of the Viral Core. J. Virol. 2015, 89 (2), 1267–1277.

50. Lee, K.; Ambrose, Z.; Martin, T. D.; Oztop, I.; Mulky, A.; Julias, J. G.; Vandegraaff, N.; Baumann, J. G.; Wang, R.; Yuen, W.; Takemura, T.; Shelton, K.; Taniuchi, I.; Li, Y.; Sodroski, J.; Littman, D. R.; Coffin, J. M.; Hughes, S. H.; Unutmaz, D.; Engelman, A.; KewalRamani, V. N., Flexible Use of Nuclear Import Pathways by HIV-1. Cell Host Microbe 2010, 7 (3), 221–233.

51. Price, A. J.; Fletcher, A. J.; Schaller, T.; Elliott, T.; Lee, K.; KewalRamani, V. N.; Chin, J. W.; Towers, G. J.; James, L. C., CPSF6 Defines a Conserved Capsid Interface that Modulates HIV-1 Replication. PLoS Path. 2012, 8 (8).

52. Luban, J.; Bossolt, K. L.; Franke, E. K.; Kalpana, G. V.; Goff, S. P., Human Immunodeficiency Virus Type 1 Gag Protein Binds to Cyclophilins A and B. Cell 1993, 73 (6), 1067–1078.

53. Hulme, A. E.; Perez, O.; Hope, T. J., Complementary assays reveal a relationship between HIV-1 uncoating and reverse transcription. P Natl Acad Sci USA 2011, 108 (24), 9975–9980.

54. Marquez, C. L.; Lau, D.; Walsh, J.; Shah, V.; McGuinness, C.; Wong, A.; Aggarwal, A.; Parker, M. W.; Jacques, D. A.; Turville, S.; Bocking, T., Kinetics of HIV-1 capsid uncoating revealed by single-molecule analysis. Elife 2018, 7.

55. Kelly, B. N.; Kyere, S.; Kinde, I.; Tang, C.; Howard, B. R.; Robinson, H.; Sundquist, W. I.; Summers, M. F.; Hill, C. P., Structure of the antiviral assembly inhibitor CAP-1 complex with the HIV-1CA protein. J. Mol. Biol. 2007, 373 (2), 355–366.

56. Sticht, J.; Humbert, M.; Findlow, S.; Bodem, J.; Muller, M.; Dietrich, U.; Werner, J.; Krausslich, H. G., A peptide inhibitor of HIV-1 assembly in vitro. Nat. Struct. Mol. Biol. 2005, 12 (8), 671–677.

57. Schneider, T.; Stoll, E., Molecular-dynamics study of a three-diemensional one-component model for distortive phase transitions. Phys Rev B 1978, 17, 1302–1322.

58. Grime, J. M. A.; Voth, G. A., Highly Scalable and Memory Efficient Ultra-Coarse-Grained Molecular Dynamics Simulations. J Chem Theory Comput 2014, 10 (1), 423–431.

